# Histone H4K20 methylation synchronizes cytoskeletal dynamics with cell cycle phases during epidermal differentiation

**DOI:** 10.1101/2020.12.01.404053

**Authors:** Alessandro Angerilli, Janet Tait, Julian Berges, Irina Shcherbakova, Tamas Schauer, Pawel Smialowski, Ohnmar Hsam, Edith Mentele, Dario Nicetto, Ralph A.W. Rupp

## Abstract

Histone tails are subject to various post-translational modifications, which play a fundamental role in altering chromatin accessibility. Although they are thought to regulate progression through development, the impact of the most abundant histone modification in vertebrates, i.e., histone H4 lysine 20 dimethylation (H4K20m2), has remained largely elusive. H4K20m2 arises from sequential methylation of new, unmodified histone H4 proteins, incorporated into chromatin during DNA replication, by the mono-methylating enzyme PR-SET7/KMT5A during G2/M phases, followed by conversion to the dimethylated state by SUV4-20H1 enzymes in the following G1/G0 phase. To address its function, we have blocked the deposition of this mark by depleting Xenopus embryos of SUV4-20H1/H2 methyltransferases, which convert H4K20 monomethylated to di- and tri-methylated states, respectively In the frog larval epidermis this results in a severe loss of cilia in multiciliated cells (MCC), a key component of all mucociliary epithelia. MCC precursor cells are correctly specified and amplify centrioles, but ultimately fail in ciliogenesis due to perturbation of cytoplasmic processes. Genome wide transcriptome profiling reveals that SUV4-20H1/H2 depleted ectodermal Animal Cap explants preferentially down-regulate the expression of several hundred cytoskeleton and cilium related genes as a consequence of persistent H4K20 monomethyl marks on postmitotic chromatin. Further analysis demonstrated that knockdown of SUV4-20H1 alone is sufficient to generate the MCC phenotype and that overexpression of the H4K20m1-specific histone demethylase PHF8 rescues the ciliogenic defect in significant, although partial, manner. Taken together, this indicates that the conversion of H4K20m1 to H4K20m2 by SUV4-20H1 is critical to synchronize cytoskeletal dynamics in concert with the cell cycle.

## INTRODUCTION

Methylation of histone residues is a major means by which epigenetic regulation of gene expression is achieved. For example, the main repressive epigenetic marks in higher eukaryotes are the trimethylation of histone H3K9, H3K27, and H4K20 (Hyun *et al*, 2017; Saksouk *et al*, 2014). Usually when a cell enters S-phase, the epigenetic information contained in parental histones is partitioned to the newly duplicated DNA strands, and is effectively diluted through incorporation of via partitioning of old histones and adequate modification of newly incorporated histones (Alabert *et al*, 2015; Petryk *et al*, 2018). This is not the case for histone H4K20 mono and di-methylation. In fact, these epigenetic marks are uniquely written in a cell cycle-dependent manner (Beck *et al*, 2012b; Pesavento *et al*, 2008). During the S-phase of the cell cycle newly synthesized, unmodified histones are inserted into the nascent chromatin (Scharf *et al*, 2009). These histones are monomethylated on H4K20 in a genome-wide fashion by Pr-Set7/Set8, a histone methyltransferase, whose activity is restricted to the G2-M phases of the cell cycle and undergoes proteolytic degradation in G1 (Beck *et al*, 2012a). After the anaphase, cells enter the G1 phase, when SUV4-20H1 and SUV4-20H2 enzymes convert this modification to the di and tri-methylated state of H4K20, respectively (Schotta *et al*, 2004). In mice, SUV4-20H1 generates H4K20m2 globally using the monomethylated residue as substrate (Pannetier *et al*, 2008). In this way during the G1/G0 phase H4K20me1 disappears from the chromatin with exception of some loci where it is shielded by unknown factors (Jorgensen *et al*, 2013). Remarkably, in both Xenopus tadpoles and mouse cells H4K20me2 covers ≥ 80% of the H4 molecules and it is considered to be the most abundant histone modification in vertebrates (Pokrovsky *et al*, 2020; Popov *et al*, 2017).

H4K20 methylation states also have distinct functional connotations. In mammals, H4K20me1 has demonstrated roles in replication origin firing (Shoaib *et al*, 2018), guaranteeing genome integrity (Oda *et al*, 2009), and chromosome condensation (Beck *et al.*, 2012a). H4K20me1 contributes to the downregulation of X-linked genes during dosage compensation in C. elegans (Brej et al., 2018), and in mammalian cells H4K20me1 down-regulates genes associated with cytoskeleton organization and cell adhesion. It has also been shown that converting H4K20me1 into H4K20me2 during the G2/M phase of the cell cycle leads to strong mitotic defects (Julien & Herr, 2004). However, due to its ubiquity, the specific functions of H4K20me2 remain elusive (Jorgensen *et al.*, 2013). H4K20me3 is a transcriptional repressor. It primarily localizes to centromeres, telomeres and repetitive DNA elements, and it participates in heterochromatin formation by recruiting factors such as cohesins (Hahn *et al*, 2013).

Here we report that depletion of SUV4-20H enzymes during Xenopus development strongly affects the formation of ciliary axonemes, the major specialized cytoskeleton structure of multiciliated cells (MCCs) on the embryonic epidermis. By taking advantage of embryonic epidermal organoids (Animal Caps) we demonstrate that knocking down the SUV4-20H enzymes negatively affects the expression of hundreds of cilia and cytoskeleton genes with striking specificity. This study shows that such repression is elicited by the high abundance of H4K20me1 on postmitotic chromatin. In wildtype cells this repressive effect is neutralized by conversion to H4K20me2 status through SUV4-20H1. Taken together our data strongly support a model in which H4K20 mono- and di-methylated states are used to equilibrate the cytoskeleton dynamics of proliferating, undifferentiated cells with those of post-mitotic differentiating cells.

## RESULTS

### SUV4-20h enzymes are required for differentiation of multiciliated cells in Xenopus larval epidermis

In order to study the function of H4K20 methylation in vivo, we induced protein knockdown (KD) of SUV4-20H enzymes via radial injection of translation blocking antisense Morpholino oligonucleotides (Mo) with established specificity and effectiveness (Nicetto *et al*, 2013), and investigated the consequences on H4K20 methyl marks (Figure 1A). SUV4-20H2 KD had a stronger effect on H4K20me3 levels than SUV4-20H1 KD, similar to what is observed in mouse, but only the double KD of both enzymes abolished the H4K20me3 mark. This indicates that the function of the two enzymes is partially redundant. Interestingly, upon individual or double KD of the SUV4-20H enzymes we observed a very strong enrichment for H4K20me1, indicating that in Xenopus this histone mark also serves as a substrate for SUV4-20H methyltransferases.

**Figure 1:**
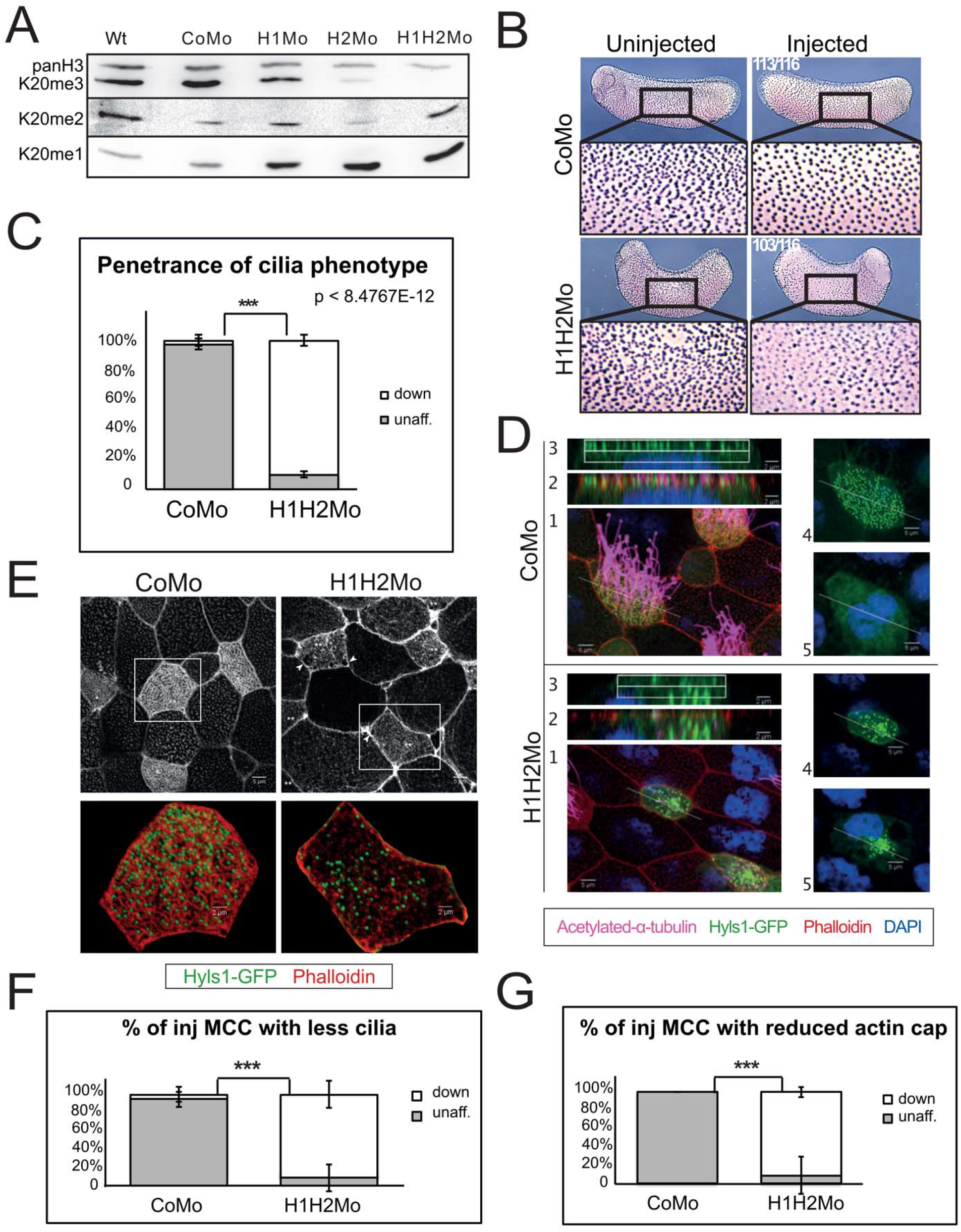
SUV4-20H1/H2 enzymes are required for ciliogenesis. A) Western Blot analysis of H4K20 methylation states upon KD of the SUV4-20H1/H2 enzymes, individually (H1Mo, H2Mo) or in concert (H1H2Mo) (n=2 biological replicates). B) Immunocytochemistry against acetylated-alpha tubulin on embryos that were half injected with either the control or both H1/H2 Mo (n=6 biol. repl.). C) Penetrance of the phenotype presented in panel B. Panels D-G: Confocal analysis of multiciliated cells on the embryonic epidermis in embryos that were mosaically injected with either control morpholino (CoMo) or the H1H2 Mo. In green are shown the basal bodies, in red the actin meshwork, in magenta the ciliary axonemes and in blue the cell nuclei. Images: 1 shows a representative MCC, 2 shows overlap between the actin cap staining and the basal bodies on the apical surface, 3 shows only the basal bodies on the uppermost Z-sections, 4 shows an apical view of the basal bodies, 5 shows a deep Z-section close to the cell nucleus. E) Confocal analysis of the actin meshwork and docked basal bodies in CoMo or H1H2Mo injected MCC. Panels F & G: Quantification of the number of MCCs showing reduced cilia (as observed in panel D) and filamentous actin staining (as in panel E) following confocal analysis. We measured 144 CoMo and 163 H1H2Mo injected multiciliated cells. Error bars represent standard deviations. For cilia p<3.61522E-08 while for actin p<3.50952E-11.

We noticed that SUV4-20H1/H2 double-morphant (dMO) tadpoles failed to exhibit the typical forward-sliding movement over ground, which is caused by the synchronous and directional beating of motile cilia on the larval epidermis. By ejecting an aqueous dye close to the surface of manipulated embryos we found that this liquid flow was strongly impaired in SUV4-20H dMO embryos (Supplementary videos). A reduced flow reflects either a problem with polarization of the cilia stroke direction, which is controlled by the PCP pathway (Mitchell *et al*, 2009), or a dysfunction of the cilia tufts on the skin surface. To distinguish between these possibilities, we injected embryos unilaterally with control or suv4-20h1/h2 morpholinos and performed whole mount immunostainings against acetylated alpha-tubulin, a major component of ciliary axonemes (Figure 1B-C). The vast majority of SUV4-20H dMOs displayed clearly reduced ciliary signals on their injected side, indicative of a cell-autonomous defect. This phenotype was reproduced with a second pair of suv4-20h1/h2-specific Mos (Supplementary material, Figure S1C and D), whose target regions are non-overlapping with that of the first Mo pair (Figure S1E). In addition, cilia tufts were significantly restored by co-injection of morpholino-insensitive Xenopus suv4-20h1/h2 mRNAs, but not by co-injection of lacZ mRNA. We also noticed that overexpression of xSUV4-20H proteins alone increased the density of MCCs within the epidermis, while lacZ mRNA had no effect (Figure S1A and B).

To better elucidate the molecular features of this phenotype we performed a cell mosaic analysis by injecting a single ventro-animal blastomere at the 8 cell-stage, whose descendants become mostly epidermis, and analysed the consequences by confocal microscopy. The injected cells were lineage-traced by hyls1-GFP mRNA, which encodes a widely conserved protein stably incorporated into the outer centriolar wall (Dammermann *et al*, 2009). In this way the progeny of the injected blastomere intermingles with the surrounding wt cells and the manipulated MCCs could be identified by GFP-positive basal bodies (BB; Figure 1D and E). We then harvested the embryos at tailbud stage (NF28), i.e. after formation of the mucociliary epithelium, and stained them for acetylated alpha-tubulin (cilia), filamentous actin (cell borders/apical actin lattice), and DNA (nucleus). This experiment demonstrated that SUV4-20H1/H2 dMO MCCs are present on the surface of the embryo and produce in deep cytoplasm a large number of centrioles (maturing into ciliogenic basal bodies). This is very similar to wildtype MCCs, which are known to generate >100 centrioles at the outer nuclear membrane via the deuterosome pathway (Meunier & Azimzadeh, 2016). A more detailed comparison with control MO (CoMo) MCCs, however, revealed that the BBs in SUV4-20H1/H2 depleted MCCs tend to clump in deep cytoplasm and were delayed in transport to, and docking at the apical cell membrane (Figure 1, compare optical sections #s 2-5 in panel D). Notably, most BBs, which had arrived at the cell membrane, still failed to nucleate ciliary axonemes, detailing a nearly complete loss of cilia in many dMO MCCs. In addition, the apical actin lattice, which forms around BBs (Sedzinski *et al*, 2016), appears much less dense and uniform in SUV4-20H1/H2 dMO MCCs, compared to CoMo MCCs. Together, these observations indicate that SUV4-20H1/H2 depleted MCCs display a differentiation defect that is due to multiple, possibly linked defects in cytoskeleton structures and processes, which are prerequisite for ciliogenesis. The severe reduction in cilia numbers is sufficient to explain the reduced liquid flow observed along the embryonic epidermis.

### SUV4-20H enzymes are epistatic to the genetic program of multiciliogenesis

The multiciliogenic differentiation program begins when an epidermal stem cell gets specified by Notch/Delta signaling as a multiciliated cell precursor (Brooks & Wallingford, 2014). Notch regulates in MCC precursors the expression of multicilin/mcidas (mci), a transcriptional coactivator protein, which establishes a core MCC transcriptome of about 1000 genes with help from key downstream transcription factors, such as Foxj1 and RFX2 (Quigley & Kintner, 2017). We therefore wondered, whether the effect of SUV4-20H1/H2 depletion on ciliogenesis could be overcome by MCI overexpression. We addressed this question in mosaic embryos, co-injecting morpholinos and hyls1-GFP transcripts with synthetic mRNA of a hormone-inducible MCI*-*GR fusion construct. We administered dexamethasone at late gastrula stage (NF12, i.e., when the endogenous mci gene gets activated) and harvested the embryos at late tailbud stage (NF28). In control embryos, MCI-GR induced both de novo amplification of centriole numbers and formation of cilia even in non-ciliated goblet cells, unambiguous evidence for an MCI gain of function phenotype (Figure S2A). In SUV4-20H1/H2 dMO embryos, however, MCI-GR failed to restore the apical Actin cap as well as ciliary axonemes (Figure S2 A-C).

To exclude the possibility that MCI could be limited in its function to activate Foxj1, a transcription factor required to produce motile cilia nucleated from the mother centriole of cells (Walentek *et al*, 2015), we also overexpressed this transcription factor in the dMO condition. As shown in Figure S2, panels D-F, the enhanced levels of Foxj1 were sufficient to induce axoneme formation in CoMo goblet cells, but could not increase cilia numbers in dMO MCCs. Taken together, these results indicate that SUV4-20H1/H2 enzymes regulate multiciliogenesis upstream of, or independently from mci and foxj1. This suggests a dominant role for adequate H4K20 methylation levels in chromatin for the assembly of motile cilia tufts at the apical surface of epidermal MCCs.

### SUV4-20H1/H2 dependent transcriptional profile of Xenopus epidermis

To analyse the changes in gene expression associated with the dMO epidermal cells, we decided to take advantage of the Animal Cap (AC) organoid system. ACs are prospective ectodermal explants that recapitulate the differentiation of the embryonic epidermis (Angerilli *et al*, 2018). We injected embryos radially at the 2-cell stage with either control or suv4-20h1/h2 Mos. We then cut ACs at the blastula stage and performed RNA-Seq analysis at three key developmental stages (gastrula, neurula and tailbud; see (Angerilli *et al.*, 2018)) in three biological replicates, obtaining approximately 100 million reads per stage. For gastrula stage ACs differential gene expression analysis revealed only 30 genes to be misregulated between CoMo and dMO conditions (23 upregulated, 7 downregulated). At the tailbud stage, we observed that dMO ACs dissociated spontaneously into single cells, indicating a problem with cell adhesion. RNA profiles of these replicates clustered heterogenously in principal component analysis (data not shown), which made it difficult to assess the differentially expressed genes. Since we had found before that the onset of epidermal differentiation in ACs is reflected by major transcriptional changes at neurula stage (NF16;(Angerilli *et al.*, 2018)), we based further analysis on this timepoint. Importantly, MCCs have already been specified in the inner cell layer, and have started to amplify centrioles and to intercalate into the outer cell layer of the epidermis (Deblandre *et al*, 1999).

We found that the expression of 3686 genes (19.7% of all genes) were altered in this dataset, the majority of them (2246) being downregulated (Figure 2A and data not shown). We then performed gene ontology enrichment analysis for both transcriptional responses to the altered H4K20 methyl landscape. The group of genes that was upregulated in the Su4-20h dMO condition was enriched for biological processes involved in chromatin organization and remodeling, methylation, gene expression and various metabolic processes (Fig. 2B). In confirmation of previous results (Nicetto *et al.*, 2013), we found oct25/pou5f3.2 and oct91/pou5f3.1 among these genes. In contrast, for the cohort of genes downregulated in dMO ACs, the top twenty entries were all associated with terms connected to cilium, centrosome, microtubules and cytoskeleton (Fig. 2C). The most enriched categories relate to “cytoskeleton” and “cilium” and comprise of hundreds of genes, which are typically expressed at medium to high levels in control ACs, and many of them are downregulated in suv4-20h1/h2 depleted chromatin (Figure 2D and E).

**Figure 2:**
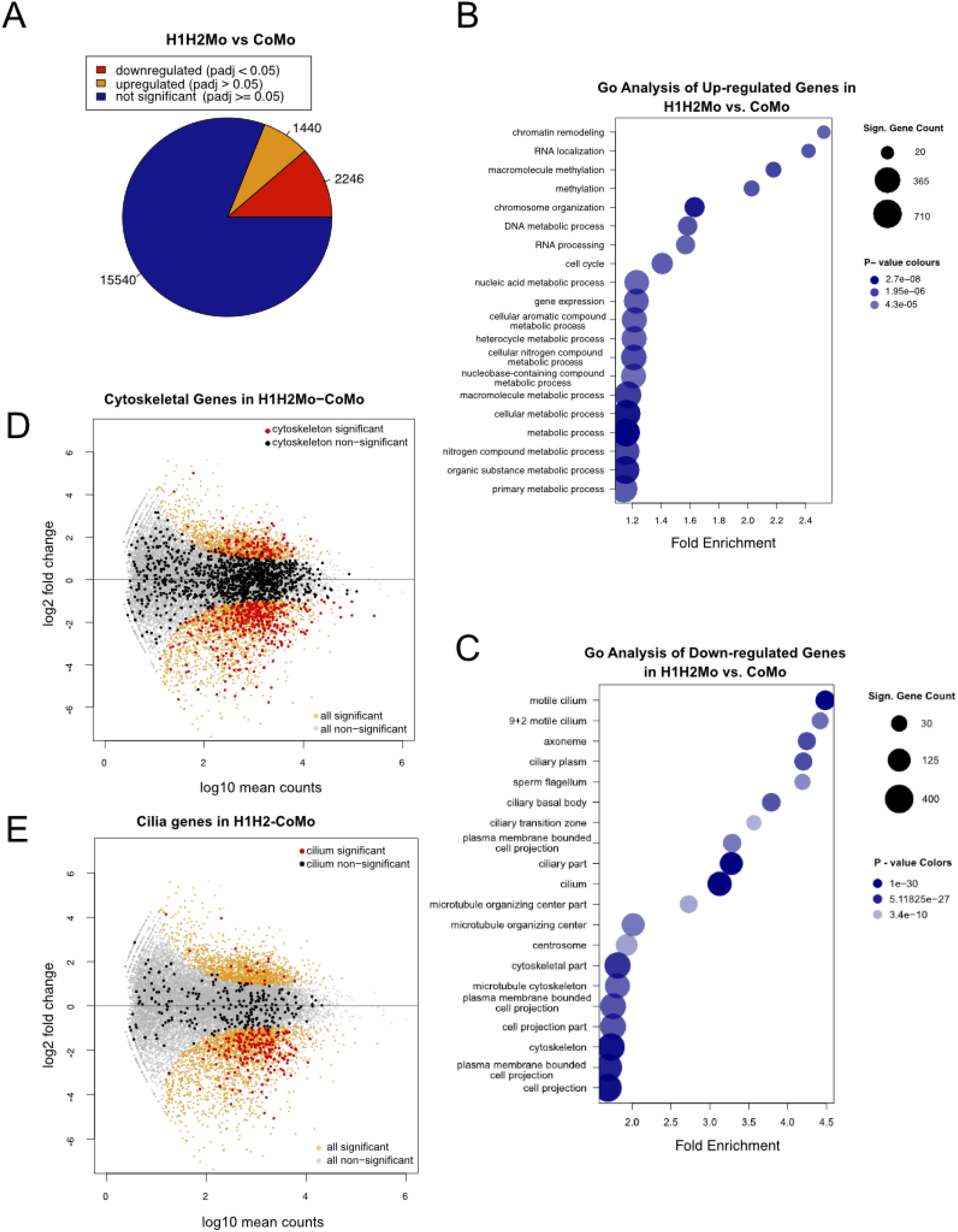
The transcriptome of SUV4-20H1/H2 depleted ACs reveals a link to cilogenesis. A) Number of misregulated genes in suv4-20h1/2 deficient NF16 ACs (significantly misregulated genes with adjusted p-value (padj) < 0.05). Gene ontology (GO) enrichment (biological process) for upregulated genes in suv4-20h1/2 morpholino injected ACs compared to control morpholino injected ACs. Bubble size represents number of significant genes per GO term and bubble colour represents p-value). C) GO enrichment for downregulated genes (cellular component) in suv4-20h1/2 morpholino injected ACs compared to control morpholino injected ACs. Bubble size represents number of significant genes per GO term and bubble colour represents p value). D, E) Gene expression in suv4-20h1/2 morpholino injected ACs compared to control morpholino injected ACs. Each dot represents a single gene. Non-significant genes are indicated in light grey and significant in orange (padj < 0.05). Cytoskeletal genes and cilium genes (as defined by R/Bioconductor package: org.Mm.eg.db version 3.8.2, mouse annotation) are indicated in black (non-significant, padj >.05), and red (significant, padj < 0.05).

Most interestingly, key transcriptional regulators of ciliogenesis were expressed either at normal (foxj1, rfx2) or slightly upregulated levels (mci) in the dMO condition (data not shown), confirming that SUV4-20H1/H2 enzymes are epistatic to these TFs. The widespread downregulation of genes associated with cytoskeleton structures and cilia provides a robust explanation for the specific MCC phenotype arising from suv4-20h1/h2 depletion. This concerted downregulation is inconsistent with the removal of repressive H4K20m3 from chromatin. Thus, another mechanism acting in a genome wide manner seems to be responsible.

### A genome-wide switch towards H4K20me1 specifically affects cytoskeleton gene expression in differentiating epidermis

The chromatin of SUV4-20H1/H2 depleted embryos is highly enriched for H4K20m1. This modification is important for mouse development (Oda *et al.*, 2009) and frequently found on promoters and coding regions of genes (Barski *et al*, 2007). Importantly, whether it represses or activates gene transcription has remained under dispute (for a discussion see (Beck *et al.*, 2012a). However, several studies on PHF8/KDM7B, a JmjC-domain containing histone demethylase with H4K20me1-specificity, have consistently revealed that increased H4K20me1 levels result in reduced gene expression (Asensio-Juan *et al*, 2017; Liu *et al*, 2010; Qi *et al*, 2010). Of particular interest to us was the finding that PHF8 knockdown in mammalian cells and primary mouse neurons caused an increase in H4K20m1 levels at coding regions, a transcriptional repression of cytoskeletal genes, impaired cell adhesion, and deficient neurite outgrowth (Asensio-Juan *et al*, 2012). Based on these phenotypic similarities to SUV4-20H1/H2 dMO Xenopus epidermis, we wondered, whether the increased H4K20me1 levels are responsible for the observed ciliogenic phenotype. To address this hypothesis, we overexpressed Phf8 in the dMO context.

We used two Phf8 cDNA clones available from commercial gene repositories: a full-length human clone (hPhf8) and a partial *Xenopus* phf8 clone (xPHF8ΔC). The latter consists of the first 264 aa residues of the xphf8 ORF, including the PHD domain known to target the protein to H3K4m3 marks at active promoters, as well as the first 66 amino acids from the 156-residue long JmjC-domain (Figure S3B). Of the critical amino acid triad His-Asp-His in the catalytic center, the truncated Xenopus protein lacks the third residue involved in Fe-coordination and, thus, is potentially compromised in its catalytic activity (Chaturvedi *et al*, 2019).

First, we injected hPHF8 mRNA, either alone or in combination with Mos, and evaluated its effect on ciliogenesis by staining half-injected embryos for acetylated-a-tubulin at tailbud stage (Figure 3A). Injection of hPHF8 alone had little to no effect on the density or size of cilia tufts, however, it restored cilia staining significantly in SUV4-20H dMOs. The injected and non-injected sides of PHF8 rescued embryos were indistinguishable in more than 40% of the inspected cases, while 90% of dMO embryos had regions devoid of cilia staining. In contrast, co-injection of the same amount of lacZ mRNA had no effect on the phenotype (Figure 3A and B). In comparable settings, xPHF8ΔC mRNA caused with high penetrance (96%) an increase in MCC numbers and ciliary staining intensity, when injected alone (Figure S3A and C). In addition, it restored ciliogenesis in a significant manner, reducing the frequency of embryos with strongly reduced cilia from 90% to 25%. In nearly half of the rescued embryos, injected and uninjected sides were indistinguishable (Figure S3A and D). In addition, we confirmed by confocal microscopy in mosaic embryos that both phf8 mRNAs were able to partially restore the assembly of axonemes and improve the density of the apical Actin meshwork, although not to the level of neighbouring wildtype MCCs (Figures 3D and S3E and F).

**Figure 3:**
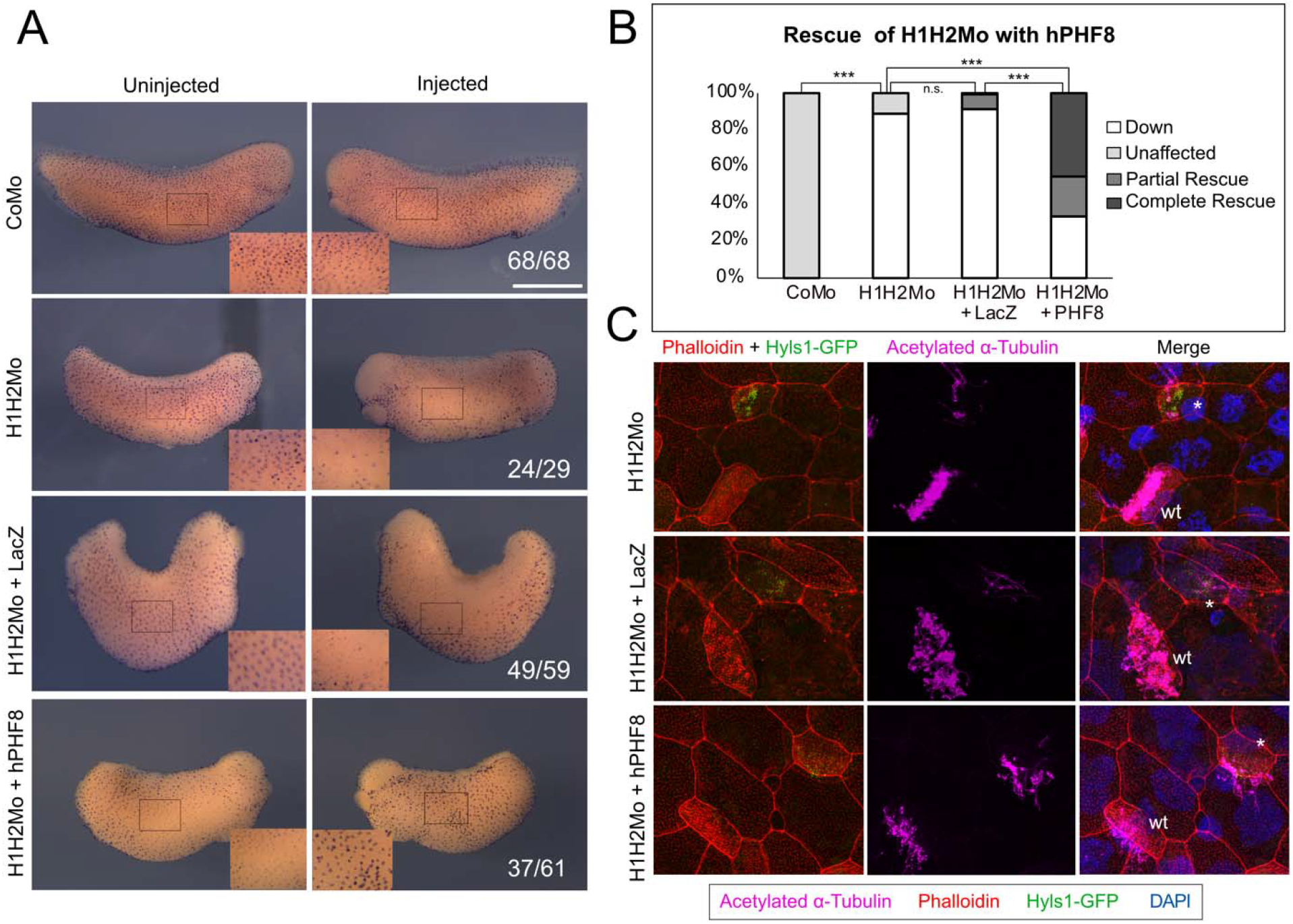
Rescue of the ciliogenic phenotype with hPHF8. A) Immunocytochemistry against acetylated alpha-tubulin of embryos that were injected with either: CoMo, H1H2Mo, H1H2Mo + 900pg of hPhf8 mRNA or H1H2Mo + 900pg of LacZ mRNA. Scale bar is 1mm. B) Quantification of cilia staining from A) in the 4 given conditions (n=3 biol. Replicates; CoMo vs. H1H2Mo: padj = 1.4E-06; H1H2Mo vs. H1H2Mo+LacZ: padj = 0.82; H1H2Mo vs. H1H2Mo+PHF8: padj = 1.132E-4; H1H2Mo+LacZ vs. H1H2Mo+PHF8: padj = 1.875E-4). D) Confocal analysis of the rescue with hPhf8 in embryos injected mosaically in 1 blastomere at the 8-cell stage. In green are shown the basal bodies, in red the actin meshwork, in magenta the ciliary axonemes and in blue are cell nuclei. * = injected MCCs, wt = wildtype MCCs.

These findings stimulated us to investigate the PHF8 dependent changes in gene expression, which accompany this partial morphological rescue. We injected embryos with hPhf8 mRNA alone or in concert with the suv4-20h1/h2 Mos (“hPHF8-rescue” condition). We also injected CoMo, and LacZ mRNA as non-specific controls. We then dissected ACs from these embryos, harvested them at neurula stage (NF16) and performed RNA-seq analysis with 3 biological replicates for each condition. The results from hPhf8, CoMo and LacZ injected explants were highly similar (data not shown). In order to stringently isolate the rescuing effect of Phf8 on the suv4-20h dMO condition, we normalized the transcriptome of hPHF8-rescue ACs by the transcriptome of Phf8-overexpressing ACs (Figure 4). Of genes that were downregulated in the initial suv4-20h1/2 condition about 80 percent had improved their expression. To correlate these changes in gene expression with the observed morphological rescue, we specifically investigated, whether transcript levels of cilium and cytoskeleton genes, which were down in the initial suv4-20h1/2 KD dataset, had improved (Figure 4B and D). Indeed, out of 183 cilium genes, 140 improved their log2-fold change (77%), while 43 were further downregulated; and out of 367 cytoskeleton genes, 294 were downregulated to a lesser extent (80%), while 73 were further downregulated (Figure 4A and C).

**Figure 4:**
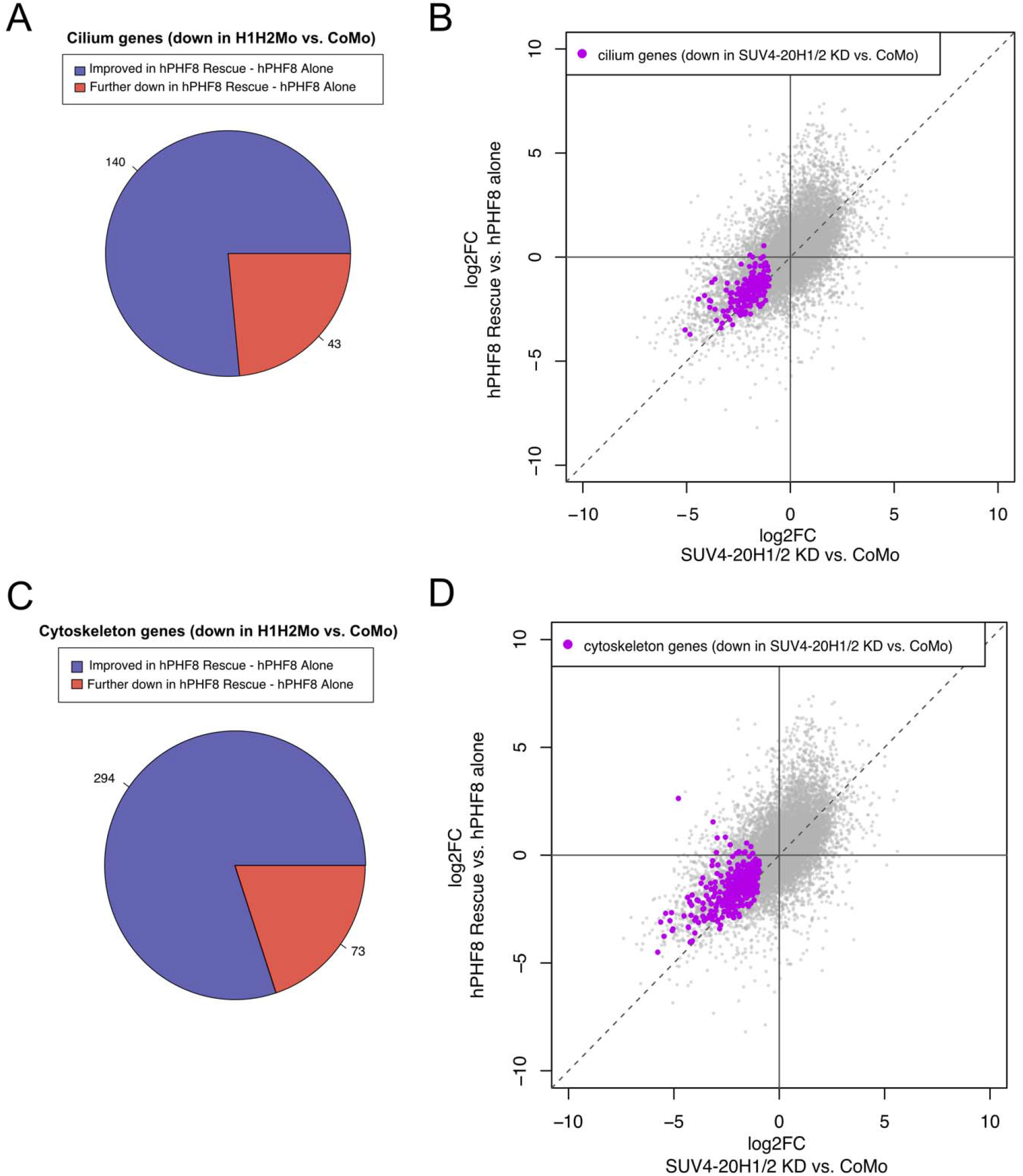
Transcriptomic analysis of hPHF8-rescued ACs. Neurula stage (NF16). (B, D) Comparison of log2 fold change between H1H2Mo vs. CoMo injected animal caps to log2 fold change of hPHF8 rescue vs. hPHF8 injected animal caps. Dashed line indicates no difference in expression between the normalized conditions. Cilia genes (B) and cytoskeletal genes (D) that were downregulated in the initial suv4-20h1/2 dMO dataset are indicated in purple. (A, Change in expression of cilia and cytoskeletal genes that were downregulated by suv4-20h1/2 knockdown upon hPHF8 rescue.

The differential gene expression analysis for the xPHF8ΔC-rescue suggested that this truncated enzyme had a stronger activity on its own than the full-length human protein (Figure S4). By comparison of the log2-fold changes, xPHF8ΔC clearly improved the expression levels of cilium and cytoskeleton genes, which were down in suv4-20h dMO condition (Figure S4D and F). This was found for 176/183 (96%) cilia genes and 344/363 (95%) cytoskeleton genes (Figure S4E and G).

All together, these results indicate that overexpression of the histone demethylase PHF8 has an influence on gene transcription in H4K20m1-enriched chromatin of suv4-20h depleted epidermal organoids. Improving the expression levels for a large majority of cilium and cytoskeleton genes that were down in SUV4-20H1/H2 depleted epidermis is compatible with the observed partial rescue of cilia tufts on the morphological level, and supports the hypothesis that the ciliogenic defect is caused through gene repression by H4K20m1.

### SUV4-20H1 is required for multiciliated cell differentiation

The suv4-20h1 gene is essential for mouse development and responsible for writing the H4K20m2 mark, while suv4-20h2 appears to be responsible for H4K20 trimethylation. Double null MEFs have lost H4K20 di- and trimethylation almost completely, without concomitant changes in acetylation or methylation levels at histone H3 lysines 9, 14, 23, 27 and 36 in bulk chromatin (Schotta *et al*, 2008). Knocking down both SUV4-20H homologs in Xenopus leads to a similar shift in H4K20 methylation states towards H4K20m1 (Figure 1A). This leads to transcriptional upregulation of some genes, which are repressed by H4K20m3, such as the mammalian OCT4-related genes oct25/pou5f3.2 and oct91/pou5f3.1 (Nicetto *et al.*, 2013). Our results from PHF8-rescue experiments in double morphant explants strongly suggest that the loss of SUV4-20H1 activity alone could already induce the ciliogenic defect during epidermal differentiation in Xenopus.

To verify this assumption, we unilaterally injected embryos with either suv4-20h1 Mo, suv4-20h2 Mo, or both and determined the effects by immunostaining for cilia tufts. As shown in Supplementary Figure 5A, the KD of SUV4-20H1 alone elicited the loss of cilia to a similar extent as the double KD of both enzymes (Figure S5B). Importantly, SUV4-20H2 depleted embryos display wildtype-like cilia tufts. In a separate experimental series, we confirmed that PHF8 partially restored ciliogenesis in SUV4-20H1 morphant explants by immunostaining (Figure 5C and D). Confocal microscopy revealed that this improves both formation of the apical actin cap and assembly of ciliary axonemes (Figure 5E). Therefore, the conversion of H4K20m1 to H4K20m2 by SUV-20H1 is essential for the differentiation of MCCs, while SUV4-20H2 is not.

**Figure 5:**
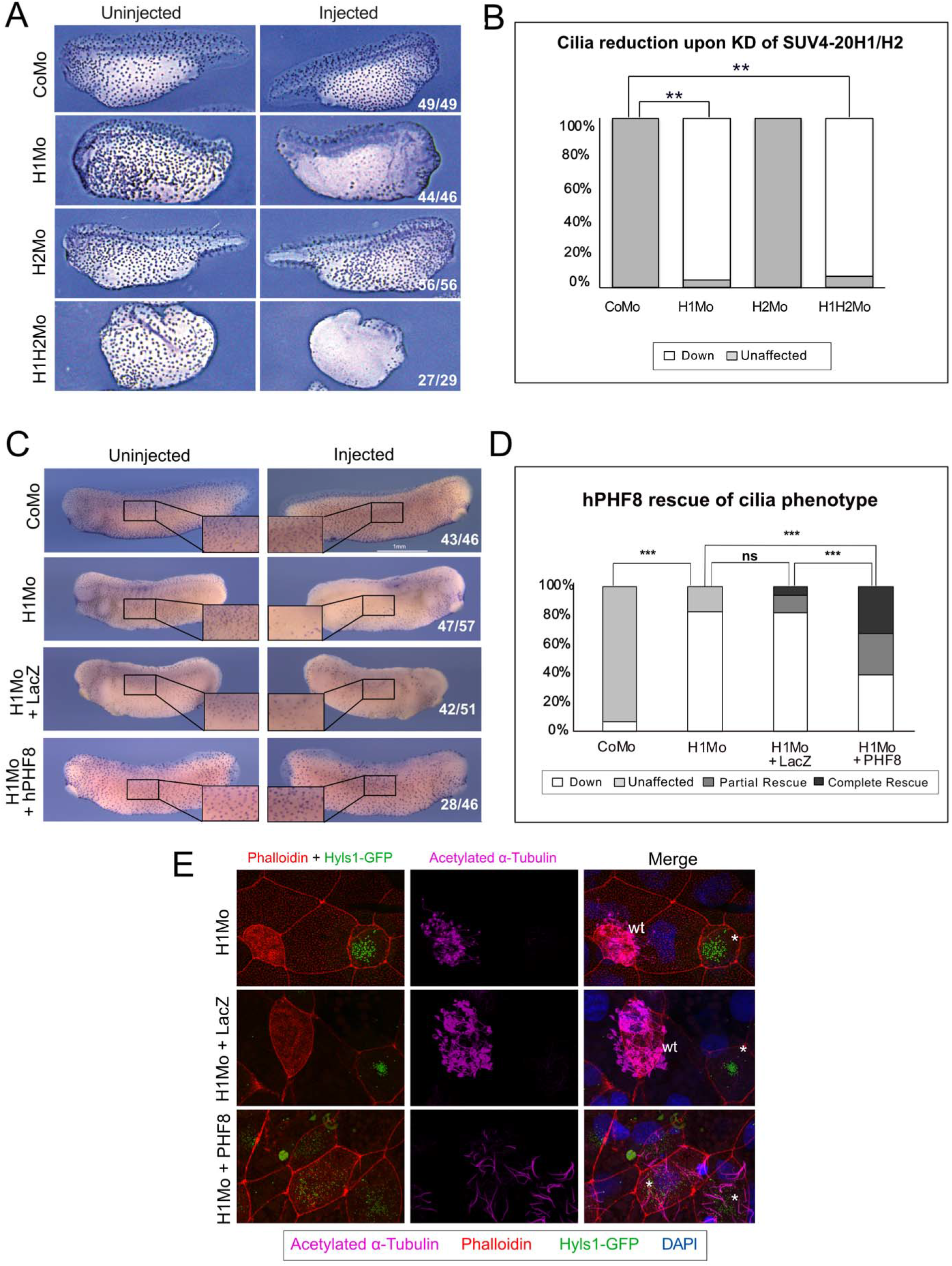
KD of SUV4-20H1 alone is sufficient to generate ciliogenic phenotype. A) Immunocytochemistry against acetylated alpha tubulin on X. tropicalis embryos that were half-injected with the suv4-20h morpholinos individually or in concert. B) Quantification of the experiment in A. For H1Mo p<0,0042, for H2Mo p<0,0015. C) Immunocytochemistry against acetylated-alpha tubulin on embryos that were half injected with CoMo, H1Mo alone, H1Mo + 900pg LacZ mRNA, or H1Mo + 900pg hPHF8 mRNA. N=4 D) Penetrance of the phenotype presented in panel C. E) Confocal analysis of multiciliated cells on the embryonic epidermis in embryos that were mosaically injected with H1Mo alone, H1Mo morpholino + 900pg LacZ mRNA, or H1Mo + 900pg hPHF8 mRNA. The basal bodies are shown in green, the actin meshwork in red, and ciliary axonemes in magenta.

## DISCUSSION

The SUV4-20H1 enzyme converts H4K20m1 to the di-methylated state and is largely responsible for making its product the most abundant histone modification in vertebrate chromatin (Jorgensen *et al.*, 2013; Pesavento *et al.*, 2008). The work presented here has elucidated an unexpected and novel function for this enzyme in developing Xenopus embryos. On the morphological level we have initially found that SUV4-20H1/H2 enzymes are needed in postmitotic MCC precursor cells at their time of differentiation to support production of a large number of motile cilia on their apical surface in the larval epidermis. These cilia tufts are needed for the generation of directional flow, a hallmark of all mucociliary epithelia (Boutin & Kodjabachian, 2019). This morphological phenotype is specific, since it was reproduced with a second pair of suv4-20h1/h2-specific Morpholino oligonucleotides. Furthermore, it was significantly rescued by injecting morpholino-insensitive suv4-20h1/h2 mRNAs, as well as by overexpression of the histone demethylase PHF8, which antagonizes the consequences of SUV4-20H1/H2 protein knockdown by removing the methyl group from hyper-accumulated H4K20m1. It should be noted that besides PHF8 there are additional enzymes, which demethylate H4K20 (Brejc *et al*, 2017; Cao *et al*, 2020). It is currently unknown, whether any of these is involved in MCC differentiation.

Differential gene expression analysis for control and SUV4-20H dMO ACs identified the underlying cause of the phenotype by revealing the concerted downregulation of about 500 genes associated with the GO terms “cilium” and “cytoskeleton”. The expression of 75% of them was improved by overexpression of PHF8 in SUV4-20H dMOs, consistent with the observed morphological rescue under this condition. Their improved expression is most likely due to changing H4K20m1 to unmethylated H4K20, which is neutral in terms of transcription. This is in agreement with a previous study, which knocked down PHF8 in mammalian cells and observed a local increase in H4K20m1 deposition at coding genes, whose expression was downregulated (Asensio-Juan *et al.*, 2017). Up to now we have not established any of the 500 genes as direct targets of H4K20m1 repression. The results from RNA profiling and cell morphological analysis, however, let us conclude that enhanced levels of H4K20m1 in chromatin is incompatible with an adequate expression of many cilium and cytoskeleton associated genes needed to support cilia tuft formation in multiciliated cells.

We subsequently found that the ciliogenic phenotype depends solely on suv4-20h1 knock down (Figure 5), and thus involves only H4K20m1 and H4K20m2 marks. This result was instrumental to formulate our model (Figure 6), which connects the ciliogenic phenotype to the cell cycle phase-specific regulation of H4K20 methylation. This model predicts that accumulation of H4K20m1 in G2-phase phase would safeguard the genome during mitosis by repressing the expression of cytoskeleton genes that interfere with either centrosome/MTOC function and/or assembly of the mitotic spindle. The incompatibility between G2/M- and G1/G0-specific gene expression programs could arise from the shared use of proteins (e.g., tubulins) and organizing hubs (e.g., centrioles/basal bodies). A major goal for the future lies in identifying the cilium/cytoskeleton genes, which need to be regulated in this manner. Once a cell exits the cell cycle, SUV4-20H1 converts H4K20m1 to H4K20m2, thereby upregulating the genes needed to form stable macromolecular assemblies in line with differentiation, such as cilia tufts, neurite outgrowths or focal adhesions. Based on this model, the ciliogenic phenotype in Xenopus MCCs is interpreted to arise from a G2-like chromatin state in a G0 cellular environment.

**Figure 6:**
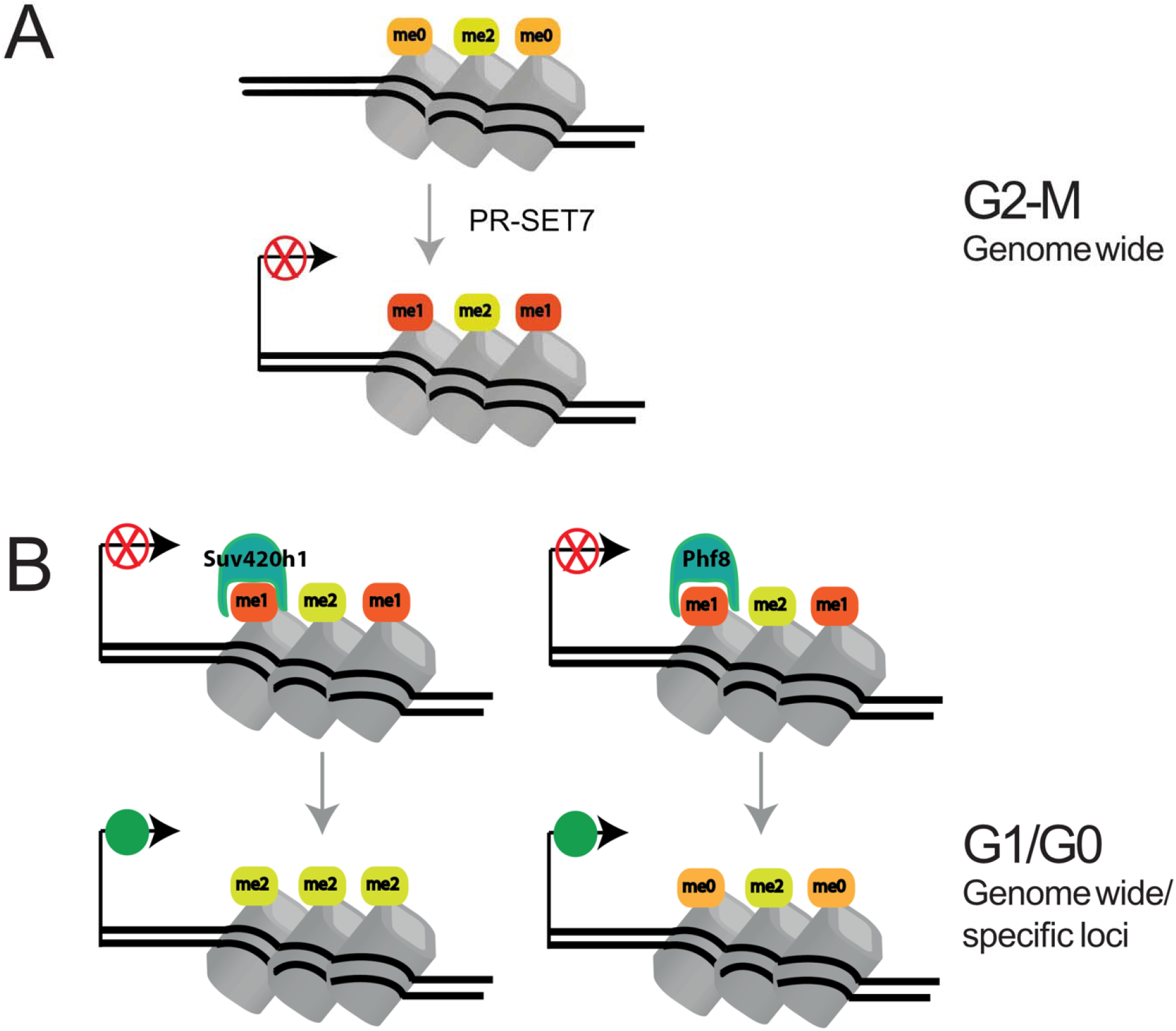
Synchronization of cytoskeletal dynamics with the cell cycle. A) During G2 and M phases of the cell-cycle, most newly synthesized H4 histones incorporated into chromatin during the preceding S-phase, become monomethylated by PR-Set7/SET8. This modification has a repressive effect on transcription, especially for cilium and cytoskeleton associated genes. B) During the following G1 and G0 phases, this repression is released by two alternative mechanisms: i) SUV4-20H1 converts H4K20me1 into H4K20me2 in a genome wide, although not quantitative fashion; ii) H4K20m1-specific demethylases such as PHF8 release repression by removing the methyl group from H4K20m1, probably in a more targeted manner.

The significance of this model is illustrated by several studies. For instance, Julien and Herr demonstrated that depletion of the abundant chromatin protein HCF-1_c_ causes a switch from H4K20m1 to H4K20m2 on mitotic chromatin, which is associated with defective chromosome alignment and segregation (Julien & Herr, 2004). Another striking example is provided by the conserved CDK1-APC/C mitotic oscillator, which at high activity levels controls entry and exit of mitosis, while at reduced levels coordinates the progression of organelle remodeling associated with motile ciliogenesis (Al Jord *et al*, 2019). Thus, postmitotic cells redeploy the mitotic oscillator to coordinate centriole amplification during multiciliogenesis (Al Jord *et al*, 2017). Although it is currently unclear, how the activity of the mitotic oscillator gets calibrated in postmitotic cells to avoid triggering mitosis, it is possible that the concerted upregulation of H4K20m1-repressed genes by SUV4-20H1 could tip the balance in favor of cytoplasmic organelle remodelling.

We assume that the fundamental mechanism we have described here has not been recognized before, since the majority of studies on histone modifications, including H4K20, were performed in proliferating cells, which are largely undifferentiated. We have detected this mechanism in the highly dynamic context of developing frog embryos, in which cells at given times stop to proliferate and switch on differentiation programs, which let them assemble diverse cytoskeletal structures depending on the cell type. It is important to recognize that the histone modification landscape of cells, which undergo this switch, is challenged by the sudden disappearance of S-phase dilution of parental histone marks through incorporation of newly synthesized, unmodified histones (Alabert *et al.*, 2015). This indeed influences epigenetic information, since histone PTMs are propagated across the cell cycle by at least two distinct kinetic modes (ibid.). In collaboration with Carsten Marr’s lab, we have recently found by computational modeling that dilution through DNA replication is sufficient to explain the measured abundance of H4K20me states in proliferating embryos, while active demethylation is needed to shape histone methylation levels in G1-arrested embryos (Schuh *et al*, 2020). We therefore believe that for a deeper understanding of histone PTMs and their impact on cellular morphology, it will be necessary to investigate the chromatin landscape in differentiated, non-proliferating cells, which after all constitute the majority of the human body.

On the organismal level, specialized cytoskeletal structures play a role in the etiology of many human diseases. For example, defects in motile cilia formation are accountable for polycystic kidney disease, Meckel-Gruber syndrome or Leber’s congenital amaurosis (Waters & Beales, 2011). Moreover cilia are essential for the mucociliary clearance in the lung and thus play a pivotal role in several diseases where clearance is impaired, such as cystic fibrosis, asthma or chronic bronchitis (Tilley *et al*, 2015). Our data show that it is possible to manipulate the amount of cilia in both directions by acting on the chromatin enzymes that regulate the abundance of H4K20 methylation. In fact, these enzymes are druggable and small molecule inhibitors for SUV4-20H enzymes and JmjC-domain containing proteins like PHF8 are available (Bromberg *et al*, 2017; Schiller *et al*, 2014). We suggest that manipulating H4K20 methylation states could have a therapeutic potential for diseases of unclear etiology that affect tubulin- or actin-based cytoskeleton structures.

## ACKNOWLEDGEMENTS

We thank Daniil Pokrovsky for helpful comments on the manuscript and its original submission; Barbara Hölscher for exceptional embryo stainings; Drs. Hiroshi Kimura, Chris Kintner, Gunnar Schotta, Axel Schweickert, and Peter Walentek for their kind gift of antibodies and recombinant plasmids; Dr. Andreas Thomae and the Core Facility of Bioimaging of the Biomedical Center for instructions and technical support in confocal imaging. This work was funded by the Deutsche Forschungsgemeinschaft (DFG, German Research Foundation) – Project-ID 213249687 – SFB 1064 (Project A12).

## MATERIALS AND METHODS

### Ethics statement

Xenopus experiments adhere to the protocol on the protection and Welfare of Animals of the European Commission and are approved by the local Animal Care authorities.

### Embryo handling and AC preparation

X. laevis embryos were handled and fertilized in vitro using standard procedures. *X. tropicalis* embryos were handled and in vitro fertilized as described (Showell & Conlon, 2009). Embryos were injected with volumes up to 10nl. 2-cell stage injected embryos were co-injected with Alexa Fluor-488 Dextran (Invitrogen) as a lineage tracer and sorted by left or right-side injection. Whole embryos were staged according to the Normal Table (Nieuwkoop & Faber, 1967). X. tropicalis animal caps (ACs) were manually dissected and staged based on sibling embryos.

### Expression constructs and morpholino oligonucleotides

Full length human phf8 cDNA in pCMV-SPORT6 and truncated xphf8 (xphf8ΔC) in pCS108 were obtained from Dharmacon™/Horizon Discovery (Cambridge UK). The following published plasmid constructs were used: Mci-HGR in pCS2+ was kindly provided by P. Walentek. Foxj1 in PCS2+ was kindly provided by A. Schweikert. Synthetic mRNAs were injected into embryos at the 2- or 8-cell stage.

Translation-blocking Morpholino oligonucleotides (Mo) directed against Xenopus suv4-20h1 (X. laevis, X. tropicalis suv4-20h1 5’-ggattcgcccaaccacttcatgcca-3’) and suv4-20h2 (X. laevis suv4-20h2: 5’-ttgccgtcaaccgatttgaacccat-3’, X. tropicalis suv4-20h2: 5’-ccgtcaagcgatttgaacccatagt-3’) and standard control Mo (5’-cctcttacctcagttacaatttata-3’) were obtained from Gene Tools LLC. *X. laevis* embryos were injected with 30-40ng of each Mo per blastomere into the animal pole of one blastomere at the 2-cell stage. For confocal analysis, X. laevis embryos were injected at the 8-cell stage in one dorsal blastomere with 5ng of each morpholino in 2.5nl (control morpholino=10ng). X. tropicalis embryos were injected with 20ng of each morpholino. Rescue experiments were performed by co-injecting suv4-20h1/2 Mos with 900pg of hphf8 mRNA, 500pg of xphf8ΔC mRNA, and equivalent amounts of lacZ mRNA.

### SDS-PAGE and western blot analysis

For Western Blot analysis the following primary antibodies were used: H4K20me1 1:2500 dilution kindly provided by H. Kimura (Kimura *et al*, 2015), H4K20me2 1:500 kindly provided by G. Schotta (Schotta *et al.*, 2008), H4K20me3 1:1000 (Abcam Ab9053), panH3 1:10000 (Abcam Ab1791), Anti-mouse 1:2500 (Jackson Immunoresearch 115-035-003), Anti-rabbit 1:5000 (Jackson Immunoresearch 1:10000).

### Immunocytochemistry

ICC was performed as described (Robinson & Guille, 1999). For confocal analysis standard methanol step was avoided. Embryos were stained with: 1:500 monoclonal anti-acetylated tubulin antibody Sigma-Aldrich (T6793), 25 μM DAPI, 0.33 μM phalloidin-Alexa 555 Thermo-Fischer Scientific (A34055). Depending on the analysis, embryos were incubated with secondary Goat anti-mouse conjugated to Alexa 647 (Thermo-Fischer Scientific A21236) or alkaline phosphatase-fused secondary Sheep anti-mouse (Chemicon AP303A). antibodies.

### Statistical Analysis

ICC results from knockdown experiments were analysis using a two-tailed Student’s t-test. Results from rescue experiments were analysed using one-way ANOVA with post-hoc Tukey test.

### RNA library preparation and sequencing

Total RNA was isolated and purified from approximately 30 X. tropicalis ACs using TRIzol (Ambion) and phenol/chloroform extraction, followed by clean-up with RNeasy Mini-Kit (Qiagen). RNA Integrity Number (RIN) was analysed using an Agilent 2100 Bioanalyzer in order to monitor RNA quality (Schroeder *et al*, 2006). Ribosomal RNA was removed from 500ng of input RNA using either Ribo-Zero Gold rRNA Removal Kit (Human/Mouse/Rat) from Illumina or NEBNext rRNA Depletion Kit (Human/ Mouse/ Rat). Then, total stranded RNA sequencing libraries were prepared using NEBNext Ultra™ Directional RNA Library Prep Kit for Illumina following the manufacturer's instructions. Quality and size of libraries were verified using the Agilent Bioanalyzer with the Agilent DNA 100 kit. RNA libraries were multiplexed and sequenced with 50 base pair (bp), paired-end reads to a depth of 30 million reads per sample on an Illumina HiSeq4000.

### RNA-seq analysis

Sequencing reads (50bp) were mapped to the reference genome (X. tropicalis v9.1 available from Xenbase) using STAR (version 2.7.1a). Gene models were provided as a gtf file, which was converted from the Xenbase gff3 format using Cufflinks gffread (version 2.2.1). Reads were counted for each gene in the same STAR run by the quantMode GeneCounts.

Downstream analyses were carried out in R (version 3.6.1) using helper function from the HelpersforDESeq2 package (version 0.1; https://github.com/tschauer/HelpersforDESeq2). Differential analysis was performed by DESeq2 (version 1.26) on the three datasets separately. Genes with at least 1 mapped read detected in 75 percent of the samples in the given dataset were considered. Significantly differential genes were defined by an adjusted p-value cutoff of 0.05. Results were visualized as MA plots, where the x-axis indicates the log10 mean counts and the y-axis the log2 fold change (log2FC). Comparison of the independent datasets was visualized by log2FC-log2FC plots, where the shown condition was compared to its own control.

Gene ontology annotation was derived from the mouse org.Mm.eg.db package (version 3.8.2) by converting gene ids from mouse to Xenopus. Genes annotated with cellular component “cilium” or “cytoskeleton” were selected by the GO ids “GO:0005929” or “GO:0005856” as well as all their offspring terms, respectively. GO enrichment analysis was performed by the topGO package (version 2.36.0) using Fisher statistics. GO results were visualized as bubble plots, where the x-axis indicates the fold enrichment (i.e. the observed number of significant genes for the GO term over the expected number), the bubble size is proportional to the number of significant genes for the GO term and the color intensity is related to the Fisher test p-value.

## Supplementary Figures

**Figure S1:**
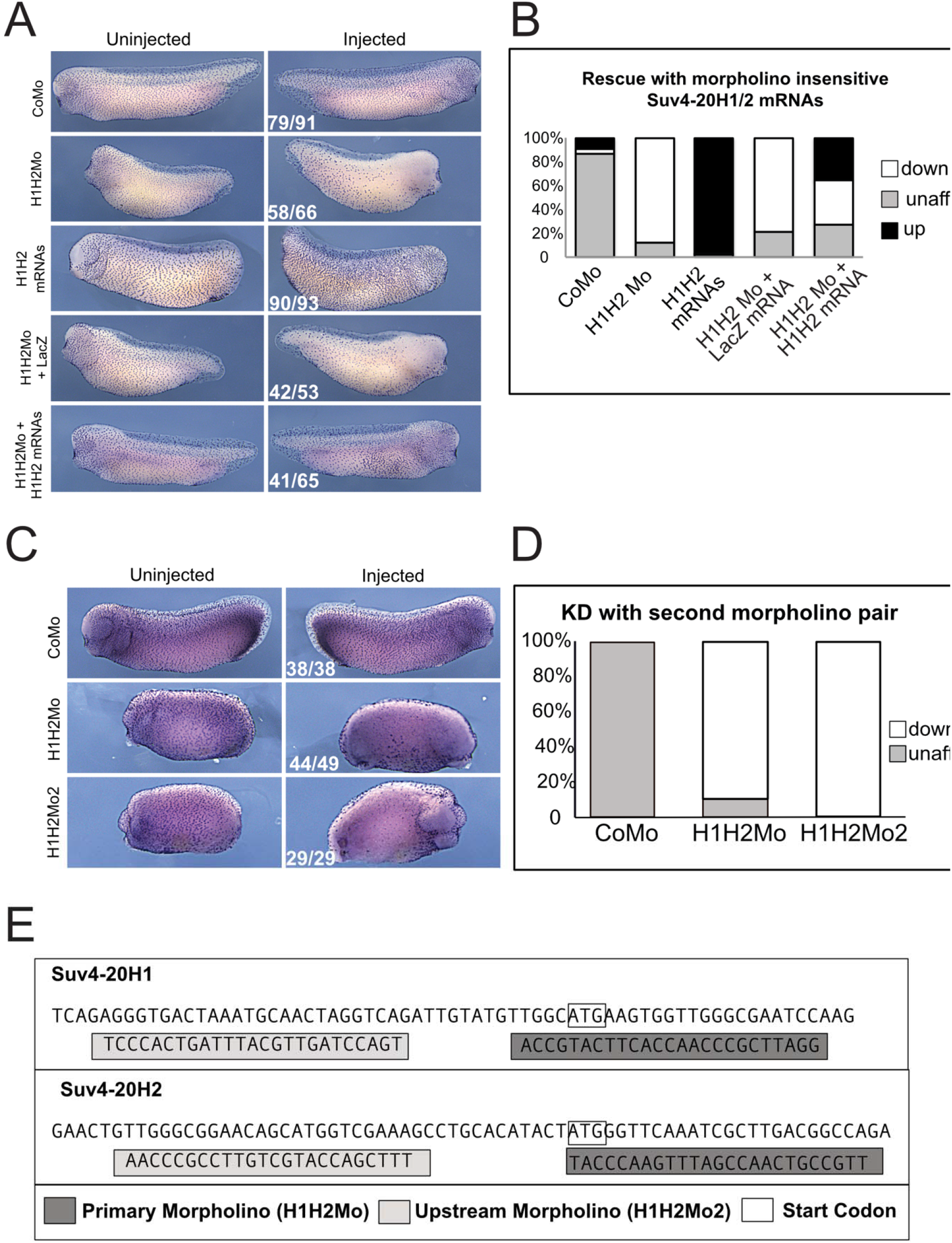
Morpholino Specificity. A) Immunocytochemistry against acetylated alpha tubulin in half injected Xenopus laevis embryos. Embryos were injected with either CoMo, the H1H2Mo, with 300pg of LacZ mRNA, or 300pg of suv4-20h1/h2 morpholino-insensitive mRNA (H1H2 mRNA) or with H1H2Mo in combination with the H1H2 mRNAs. B) Quantification of the results of the experiments in A. N=3. C) Immunocytochemistry against acetylated alpha-tubulin in embryos that were half injected with either CoMo, H1H2Mo or a second non overlapping pair of morpholinos targeting suv4-20h1/h2 (H1H2Mo2). D) shows the quantification of the experiment in C. E) Schematic representation of target regions for primary and upstream morpholinos directed against suv4-20h1 and suv4-20h2.

**Figure S2:**
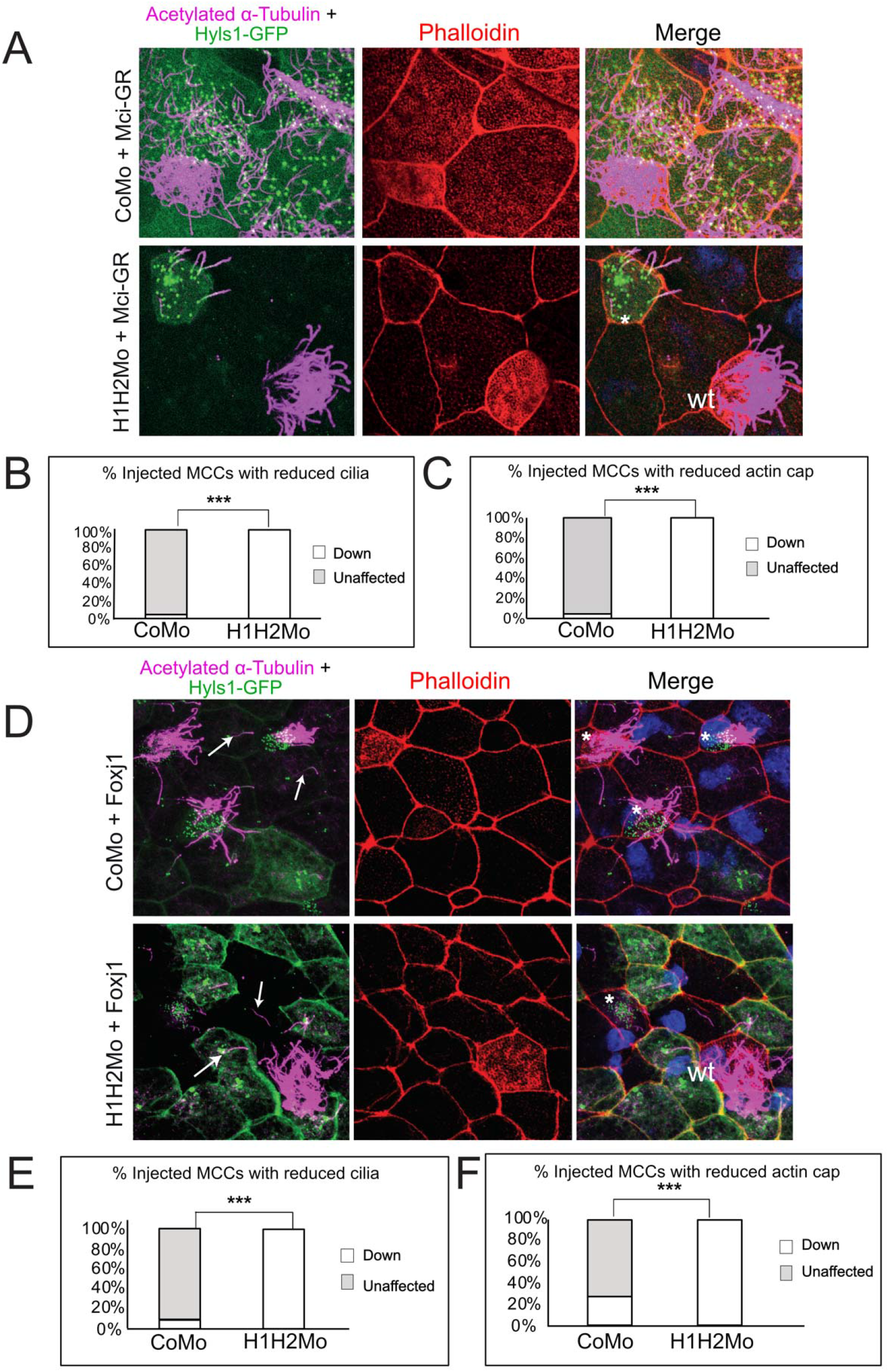
The cilia phenotype of SUV4-20H1/H2 KD embryos is not rescued by Mci/IDAS or Foxj1. A) Embryos were injected mosaically with control or suv4-20h1/2 morpholinos together with Mci-Gr mRNA. Basal bodies are shown in green, actin meshwork is shown in red, ciliary axonemes are shown in magenta, and DNA is shown in blue. * = injected MCC, wt = wildtype MCC, arrows indicate ectopic cilia. B, C) Quantification of the rescue of B) cilia (p = 3.07225E-11) and C) actin cap (p = 3.50369E-08). 8 fields of view from 8 CoMo injected embryos and 7 fields of view from 7 H1H2Mo injected embryos were scored. D) Embryos were injected mosaically with CoMo or H1H2Mo together with foxj1 mRNA. Basal bodies are shown in green, actin meshwork is shown in red, ciliary axonemes are shown in magenta, and DNA is shown in blue. E, F) Quantification of the rescue of E) cilia (p = 1.1308E-05) and F) actin cap (p = 0.00158683). 5 fields of view from 5 embryos were scored.

**Figure S3:**
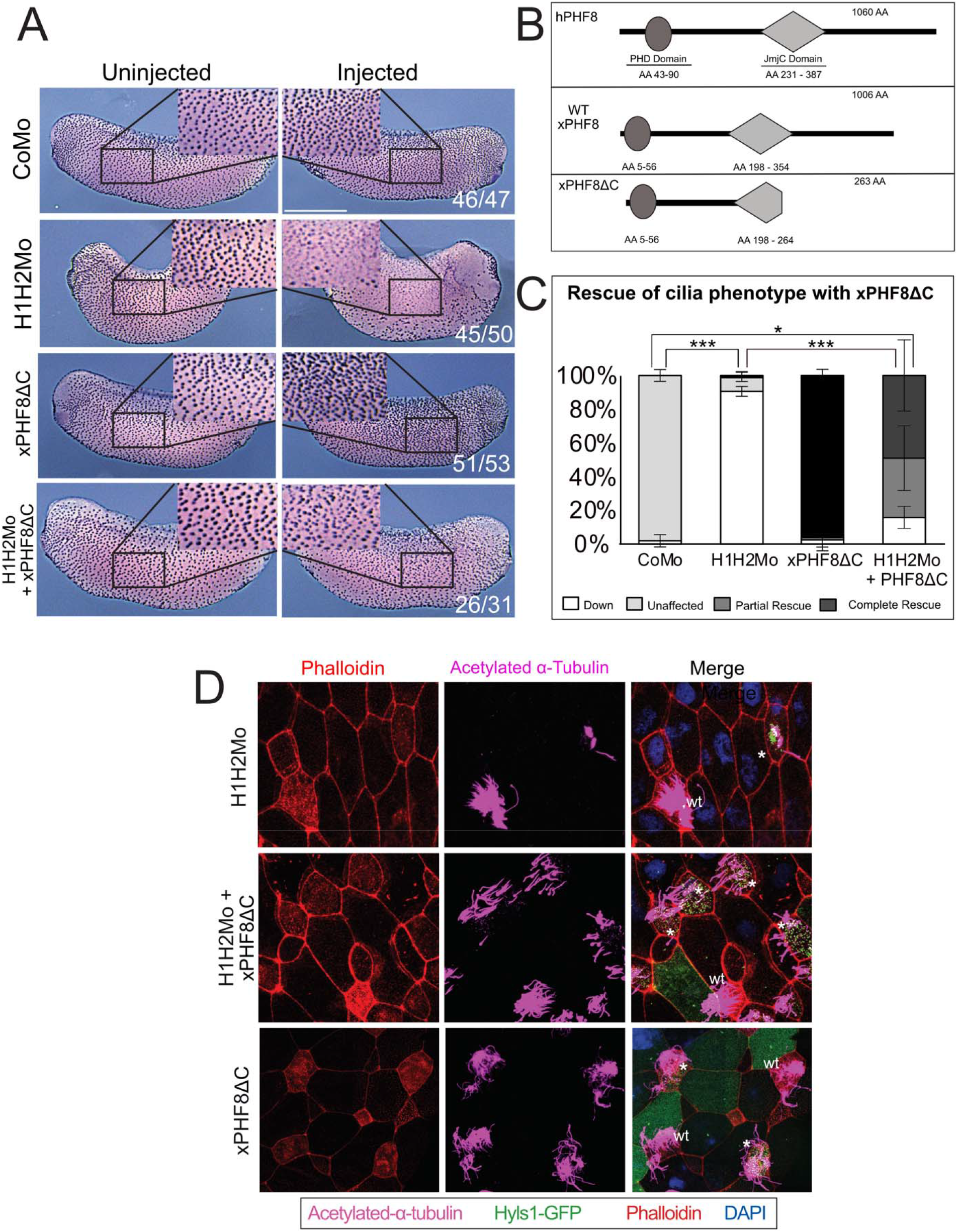
Rescue of the ciliogenic phenotype with xPHF8ΔC. A) Immunocytochemistry against acetylated alpha-tubulin of embryos that were injected with either: CoMo, H1H2Mo, xPHF8ΔC synthetic mRNA or the suv4-20h morpholinos + 300pg of xPHF8ΔC mRNA. Scale bar = 1mm, n=3. B) Schematic representation of the structure of PHF8 clones: Full length human PHF8 (hPHF8), wt Xenopus PHF8, and truncated Xenopus PHF8 (xPHF8ΔC). C) Quantification of cilia staining in panel A in the four given conditions. H1H2Mo vs. CoMo padj = 1.3E-06, H1H2Mo vs. H1H2Mo + xPHF8ΔC padj = 3.0E-06, CoMo vs. H1H2Mo + xPHF8ΔC padj = 0.0318. D) Confocal analysis of the rescue with xPHF8ΔC in embryos injected mosaically in 1 blastomere at the 8-cell stage. In green are shown the basal bodies, in red the actin meshwork, in magenta the ciliary axonemes and in blue are cell nuclei. * = injected MCC, wt = wildtype MCC.

**Figure S4:**
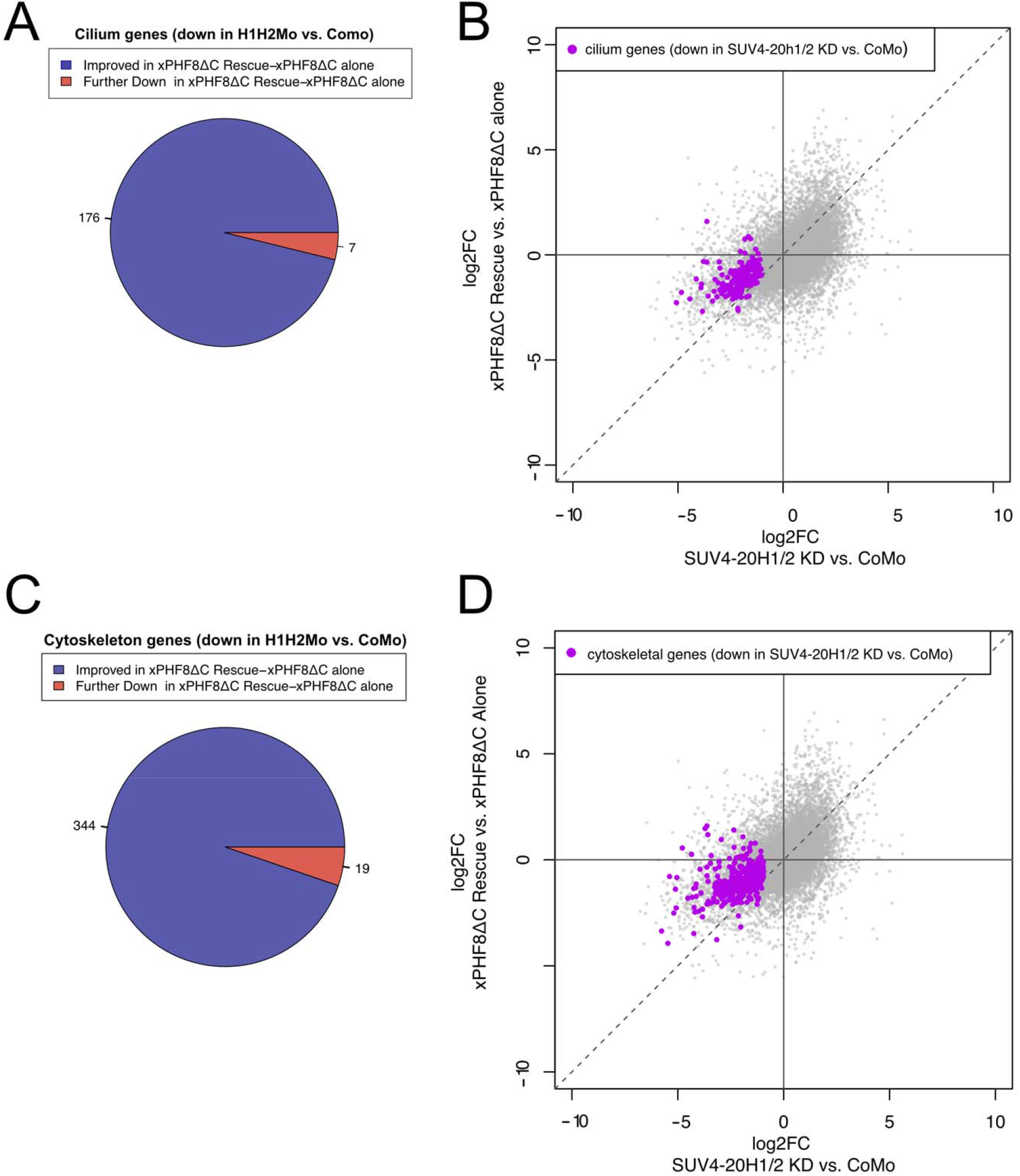
Transcriptomic analysis of xPHF8ΔC rescue ACs. Neurula stage (NF16). B, D) Comparison of log2 fold change between H1H2Mo vs. CoMo injected animal caps to log2 fold change of xRescueΔC vs. xPHF8ΔC animal caps. Cilium genes (B) and cytoskeletal genes (D) that were downregulated in the initial H1H2Mo dataset are indicated in purple. A, C) Change in expression of cilia (A) and cytoskeletal genes (C) that were downregulated by SUV4-20H1/2KD with xPHF8ΔC rescue. (A) 176 cilia genes improved their expression, while 7 were further downregulated. (C) 344 cytoskeleton genes improved their expression, while 19 were further downregulated.

